# Monitoring group activity of hamsters and mice as a novel tool to evaluate COVID-19 progression, convalescence and rVSV-ΔG-spike vaccination efficacy

**DOI:** 10.1101/2021.06.09.447687

**Authors:** Sharon Melamed, Boaz Politi, Ettie Grauer, Hagit Achdout, Moshe Aftalion, David Gur, Hadas Tamir, Yfat Yahalom-Ronen, Shlomy Maimon, Efi Yitzhak, Shay Weiss, Amir Rosner, Noam Erez, Shmuel Yitzhaki, Shmuel C Shapira, Nir Paran, Emanuelle Mamroud, Yaron Vagima, Tomer Israely

**Author notes:** Corresponding Authors: Yaron Vagima and Tomer Israely. Equal Contribution (Sharon Melamed, Boaz Politi, Ettie Grauer).

## Abstract

COVID-19 pandemic initiated a worldwide race toward the development of treatments and vaccines. Small animal models were the Syrian golden hamster and the K18-hACE2 mice infected with SARS-CoV-2 to display a disease state with some aspects of the human COVID-19. Group activity of animals in their home cage continuously monitored by the HCMS100 was used as a sensitive marker of disease, successfully detecting morbidity symptoms of SARS-CoV-2 infection in hamsters and in K18-hACE2 mice. COVID-19 convalescent hamsters re-challenged with SARS-CoV-2, exhibited minor reduction in group activity compared to naive hamsters. To evaluate rVSV-ΔG-spike vaccination efficacy against SARS-CoV-2, we used the HCMS100 to monitor group activity of hamsters in their home cage. Single-dose rVSV-ΔG-spike vaccination of immunized group showed a faster recovery compared to the non-immunized infected hamsters, substantiating the efficacy of rVSV-ΔG-spike vaccine. HCMS100 offers non-intrusive, hands-free monitoring of a number of home cages of hamsters or mice modeling COVID-19.

## Introduction

The coronavirus disease 2019 (COVID-19) pandemic initiated a worldwide race toward the development of treatments and vaccines against the emerged RNA virus Severe Acute Respiratory Syndrome Corona Virus-2 (SARS-CoV-2). Early on, the research focused on searching and defining an appropriate animal model able to represent disease symptoms and progression patterns of SARS-CoV-2 infected humans, to be used in the pre-clinical studies^1^. During earlier coronavirus sever outbreaks of Severe Acute Respiratory Syndrome (SARS-CoV-1) in 2002 and Mediterranean Respiratory Syndrome (MERS-CoV) in 2012, golden Syrian hamsters infected with various SARS-CoV strains were implemented to show viral replication and lung pathology^2^. At the present SARS-CoV-2 outbreak, hamsters were found to be an appropriate animal model of COVID-19 mainly based on the demonstration of viral replication and pathological damage at the nasal and lung tissue associated with high viral load. Their clinical symptoms however were subtle and mainly relied on weight loss^3 4^.

The specificity of the SARS-CoV-2 virus to the human angiotensin I converting enzyme 2 (hACE2) was found to be a significant barrier in developing animal models. Mice, abundantly used in pharmacological and immunological studies neither support SARS-CoV-2 infection nor exhibit any signs of morbidity following infection^1^. Consequently, a SARS-CoV-2 transgenic mouse model, known as K18-hACE2, was developed by inserting the hACE2 gene into the mouse genome^5^.

Recently, a recombinant replication competent VSV-ΔG-spike vaccine (rVSV-ΔG-spike) was developed in which the glycoprotein of VSV was replaced by the spike protein of SARS-CoV-2. A single-dose vaccination of hamsters with rVSV-ΔG-spike resulted in a rapid and potent induction of SARS-CoV-2 neutralizing antibodies. Vaccination protected hamsters against SARS-CoV-2 challenge, as demonstrated by the prevention of weight loss, and by the alleviation of the extensive tissue damage and viral loads in lungs and nasal turbinates^6^.

The large-scale use of both hamsters and mice, and the limited indications for disease manifestation or treatment efficacy required continued, high-throughput, non-intrusive method for the evaluation of disease appearance and progression and its alteration by potential vaccines and treatments. We recently introduced a method for monitoring spontaneous group activity of mice in a home cage. Validation of home cage monitoring system (HCMS100), was based on disease state induced by pulmonary exposure to LPS or influenza virus. Overall activity of mice groups tested together, demonstrated clear rate and temporal differences in activity between groups of diseased and control mice over days, in a hands-free and non-invasive fashion^7^.

Here we used the HCMS100 to detect changes in group activity of both Syrian golden hamsters and K18-hACE2 mice housed in their home cage following SARS-CoV-2 infection. We also studied the effect on group activity following re-infection in hamsters. To further corroborate rVSV-ΔG-spike vaccination efficacy against SARS-CoV-2, we used the HCMS100 to monitor vaccinated hamsters group activity and detected a faster recovery rate. This efficient, non-intrusive and safe method of monitoring continuous activity, mainly regarding BSL-3 requirements, offers a sensitive measure of disease intensity and progression as well as vaccine efficacy against SARS-CoV-2 in animal models.

## Results

Group activity of hamsters has been evaluated as a measure of animal’s health. The feasibility of monitoring the baseline activity of a group of hamsters in their home cage was examined by the use of HCMS100^7^. This system is based on a single laser beam and a detector that horizontally crosses the cage at the feeding zone and animals crossing the laser beam trigger an event recorder (Fig. 1A). Activity was clearly higher during the night hours compared to day time in these nocturnal animals as can be seen in the detailed cumulative record of night vs. day activity presented in 10 min bins over 20 days. Activity record was reset at light change (5 am and 5 pm, Fig. 1B). These data are summarized separately for day and night total activity per 20 days (Fig. 1C). The overall pattern of changes in activity is demonstrated in Fig 1D in which activity was averaged (±SEM) for each 10 min bin over the 20 days recorded. These data revealed a peak activity at approximately 1-2 h into the night cycle. Similar pattern was also observed in hamsters in a single animal testing^8^ and is different from the two-peaks pattern at light shift (day to night and night to day) seen in groups of mice^7^.

**Figure 1:**
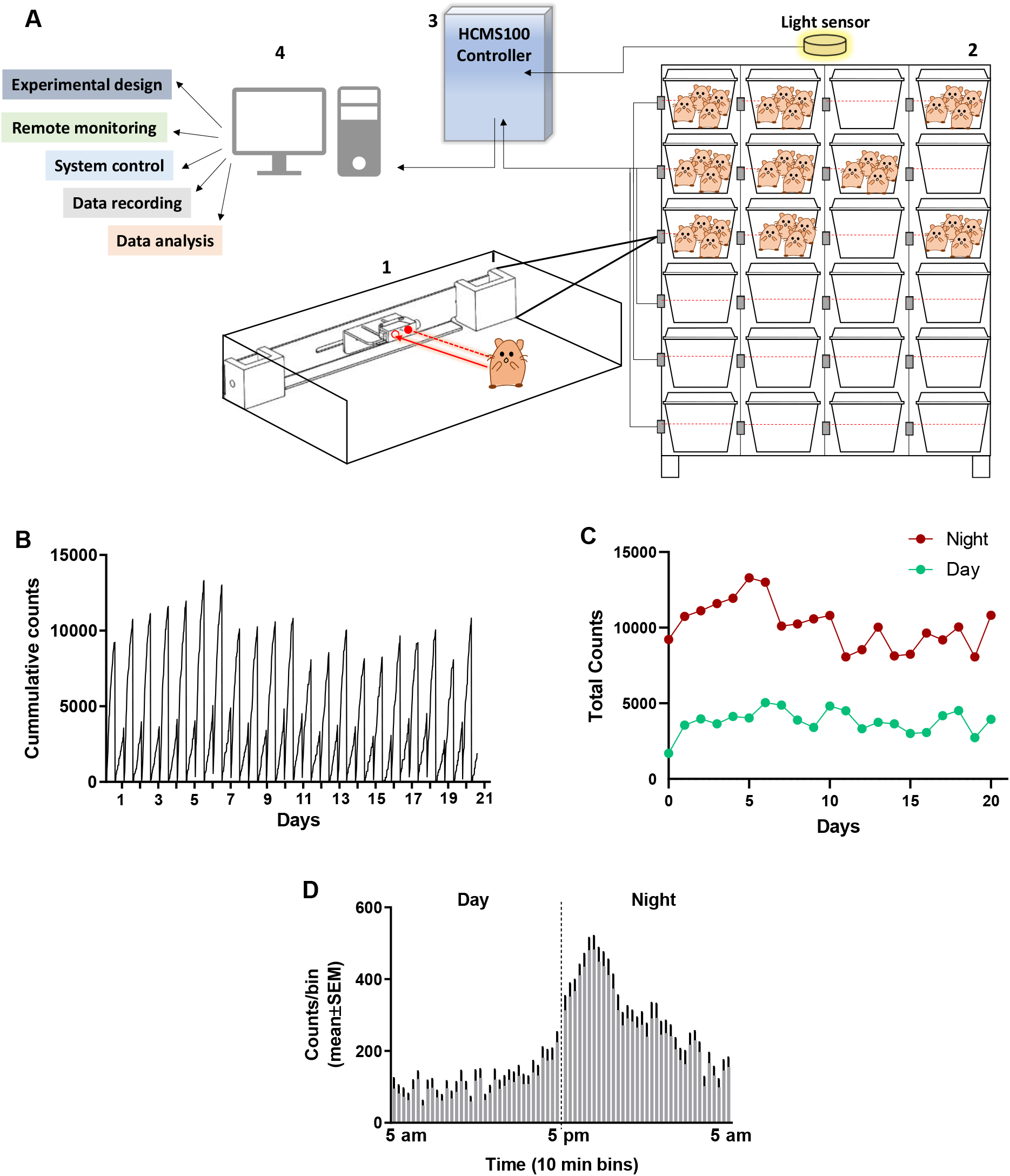
Total activity of 4 Syrian golden hamsters in their home cage recorded continuously over 20 days using the HCMS100. (A) HCMS100 schematic outline - A single retroreflective sensor is located externally to each cage at the feeding zone area (1), multiple cages with 4 hamsters/per cage, are located on a cage rack wired with an individual sensor for each cage (2). The detectors are connected to communication controller (3) with a light sensor to enable data collection and analysis of circadian activity in the cage (4). (B) Activity counts of a group of 4 hamsters in a home cage in 10 min bins with lights on from 5 am to 5 pm. Cumulative activity record resets at light change (5 am / 5 pm). (C) Total counts of activity during night time (5 pm–5 am) vs. day time (5 am– 5 pm) over 20 days. (D) Activity counts for the same group of hamsters’ averaged (±SEM) in 10 min bins over 20 days to demonstrate the overall pattern of nocturnal increase in activity, peaking approximately 1-2 h into the dark cycle.

Group activity of male and female hamsters infected intranasal (i.n.) with different doses of SARS-CoV-2 were continuously monitored by the HCMS100 (Fig. 2). Control (Mock) females were more active than males during night time with average counts of ~ 9000 counts/12h compared to ~ 5500 counts/12h counts, respectively. Two days post SARS-CoV-2 infection, both females and males showed a substantial decrease in activity that returned to normal 8 days post infection (dpi) (Fig. 2 A, D). Both females and males showed similar day time baseline activity with similar decrease in activity rates after infection. (Fig. 2 B, E). Reduced activity was accompanied by changes in weight loss (Fig. 2 C, F) in both males and females. Comparison between the two markers of disease state, namely activity and weight loss, show that changes in activity precede the changes in body weight, thus highlighting the importance of activity monitoring to evaluate COVID-19 disease progression.

**Figure 2:**
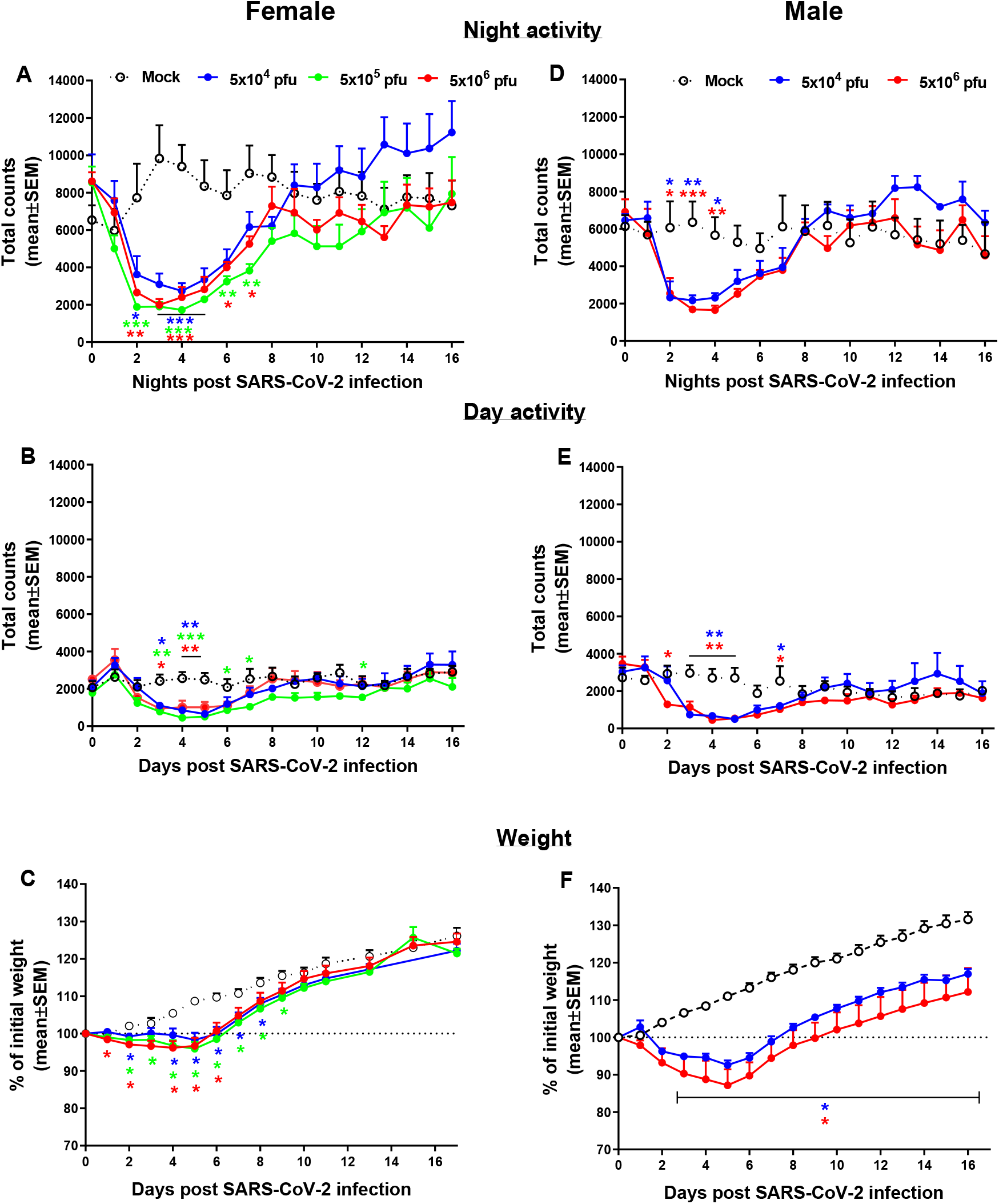
Group activity and body weight changes in SARS-CoV-2 infected hamsters. Total counts of activity during night time (5 pm – 5 am, A, D), day time (5 am – 5 pm, B, E) and weight changes (C, F) over 16 days of female (A, B and C) and males (D, E and F) hamsters infected i.n. with different doses of SARS-CoV-2. n=3-5 cages with 4 hamsters/cage. *p<0.05, **p<0.01, ***p<0.001 vs. Mock. Asterisks are color coded according to viral load as indicated.

Transgenic mice expressing the human angiotensin I-converting enzyme (ACE2) receptor driven by the cytokeratin-18 gene promoter - K18-hACE2, an additional model for the SARS-CoV-2 infection, were also tested here for changes in group activity following i.n. infection with a lethal dose of 2000 pfu of SARS-CoV-2 (Fig. 3). Infected female mice showed morbidity effect seen as a sharp decrease in both their night and day time activity 4-5 days following SARS-CoV-2 infection, accompanied by reduction in their body weight (Fig 3C). All animals died within 6-7 days following infection.

**Figure 3:**
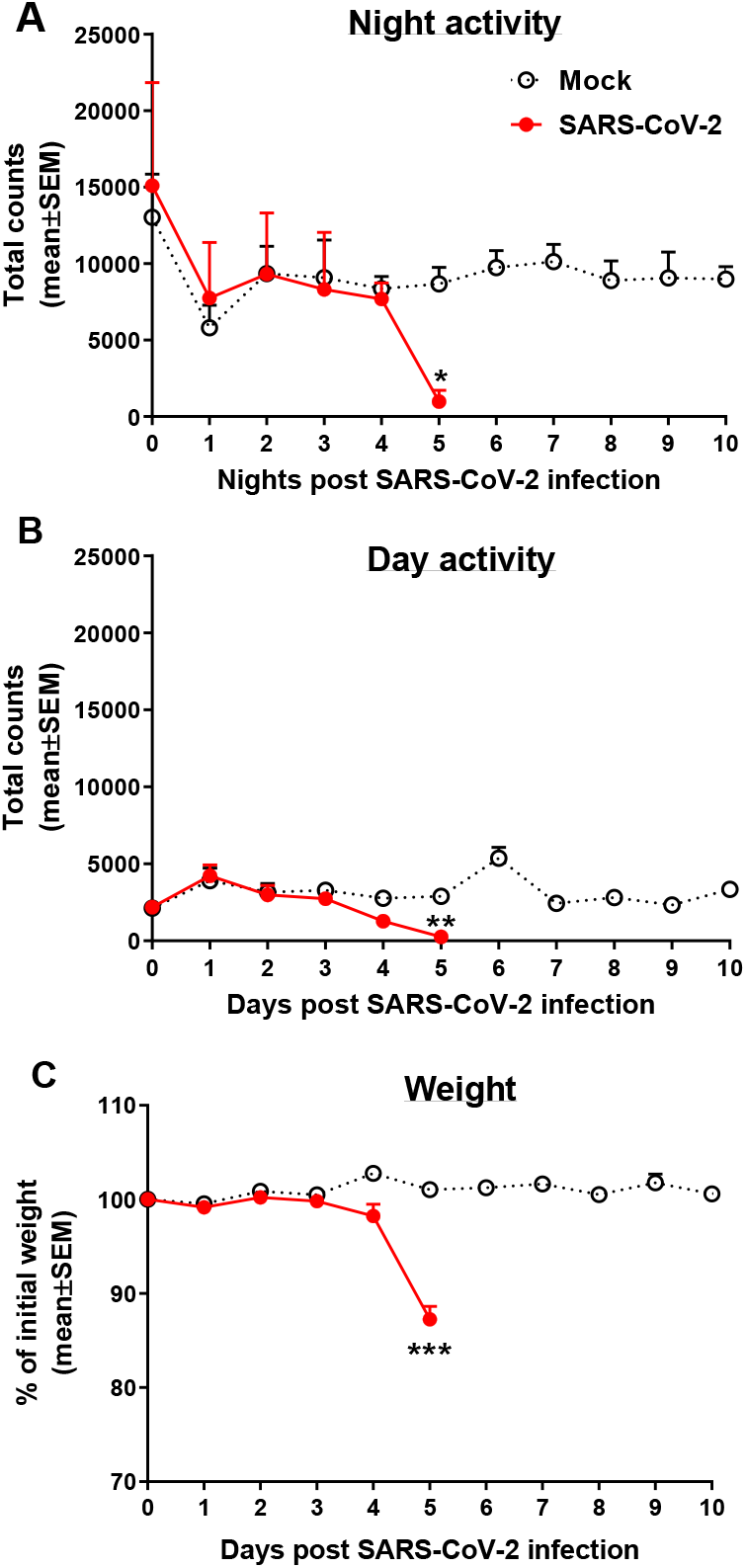
Group activity and weight changes of K18-hACE2 female mice infected with SARS-CoV-2. Total counts of activity during night time (5 pm – 5 am, A) and day time (5 am – 5 pm, B) and weight changes (C) over 10 days of mice infected i.n. with 2000 pfu of SARS-CoV-2. None of the infected animals survived the infection (death on days 6-7). Infection exposure on day 0, n=4 cages/group, 5 mice/cage. *p<0.05, **p<0.01, ***p<0.001 vs. Mock.

The use of hamsters as a model to develop treatment against SARS-CoV-2 infection was first tested for the effects of re-exposure of convalescent hamsters to SARS-CoV-2 (Fig. 4). Male hamsters initially exposed to 5×10^6^ pfu of SARS-CoV-2, showed the previously reported decrease in both day and night time activity followed by a full recovery (see Fig. 2). After 35 days following the first exposure and returning to baseline activity, the hamsters were re-infected with the same dose of 5×10^6^ pfu SARS-CoV-2 and disease progression was compared to first time infected age match hamsters. Figure 4 is a summary of the effects following re-exposure of the same animals to SARS-CoV-2, presented as percent of activity counts. The decrease in activity following the second infection (re-infection) was 40% at its peak on day 3 after re-infection compared to 70-80% decrease seen from day 2 after the first infection of naïve animals that were exposed to the same viral load.

**Figure 4:**
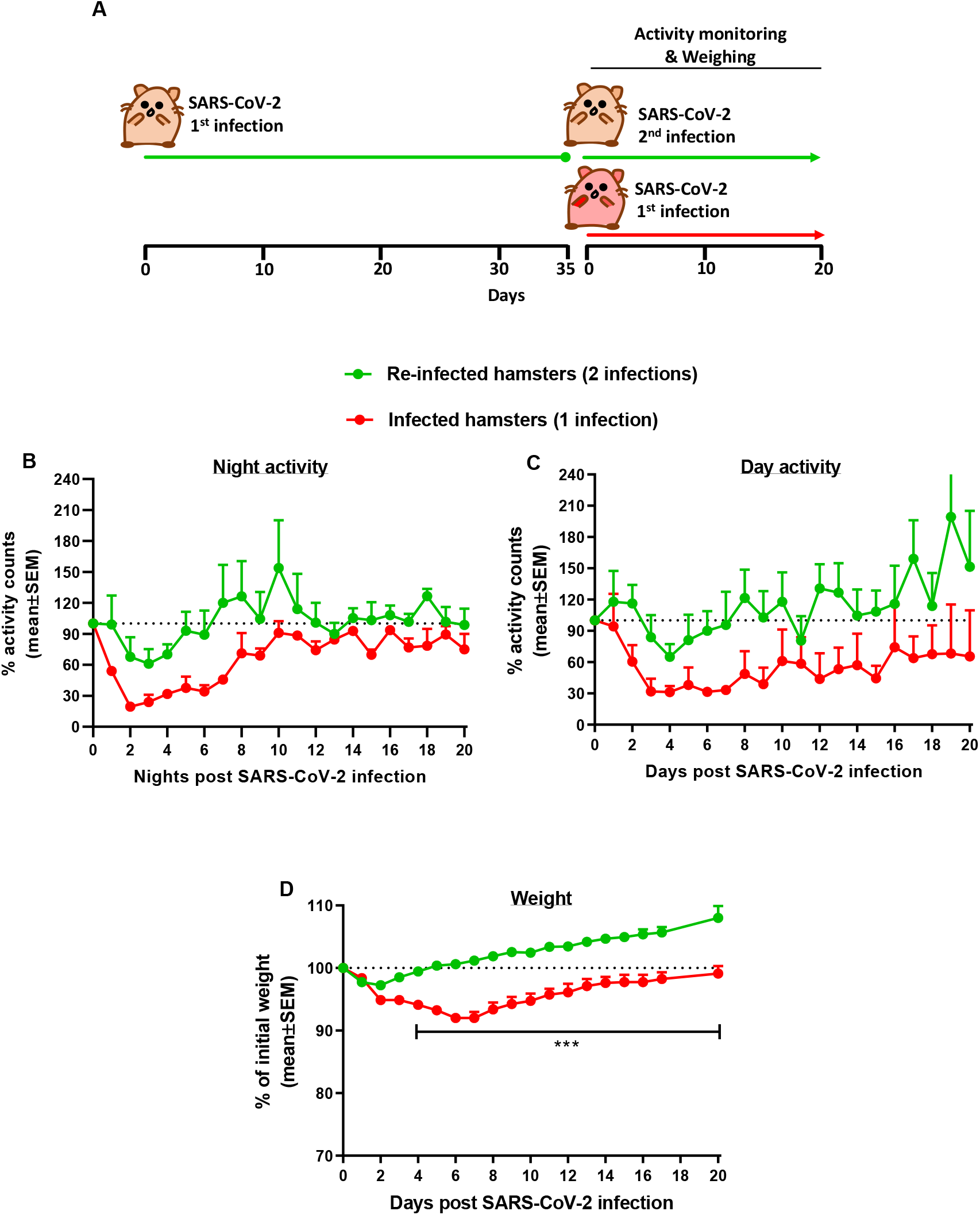
Group activity and body weight changes in male hamsters re-infected with SARS-CoV-2 after recovery. (A) Experiment outline: SARS-CoV-2 i.n. infection (5×10^6^ pfu/hamster) 35 days after the first infection with the same dose. Percent of total activity counts during night time (5 pm – 5 am, B), day time (5 am – 5 pm, C) and weight changes (D) over 20 days compared to 1^st^ time infected. n=4 cages of re-infected and 2 cages of first time infected 4 hamsters/cage. ***p<0.001

Continuous monitoring of group activity was further used to study the efficacy of rVSV-ΔG-spike vaccine as depicted in the scheme (Fig. 5A). Hamsters were immunized with 1×10^6^ pfu of rVSV-ΔG-spike and monitored for 22 days. We observed normal circadian activity following vaccination similar to that of naïve hamsters (data not shown). On day 23 both immunized and non-immunized groups were infected with 5×10^6^ pfu of SARS-CoV-2 and the previously reported night time decreased activity was clearly seen. Recovery rate was determined here as percent of activity at disease state seen 2 days after infection with 5×10^6^ pfu of SARS-CoV-2. Recovery of hamsters pre-immunized with rVSV-ΔG-spike vaccine was seen as a faster and significantly higher group activity compared to non-immunized hamsters (Fig. 5 B).

**Figure 5:**
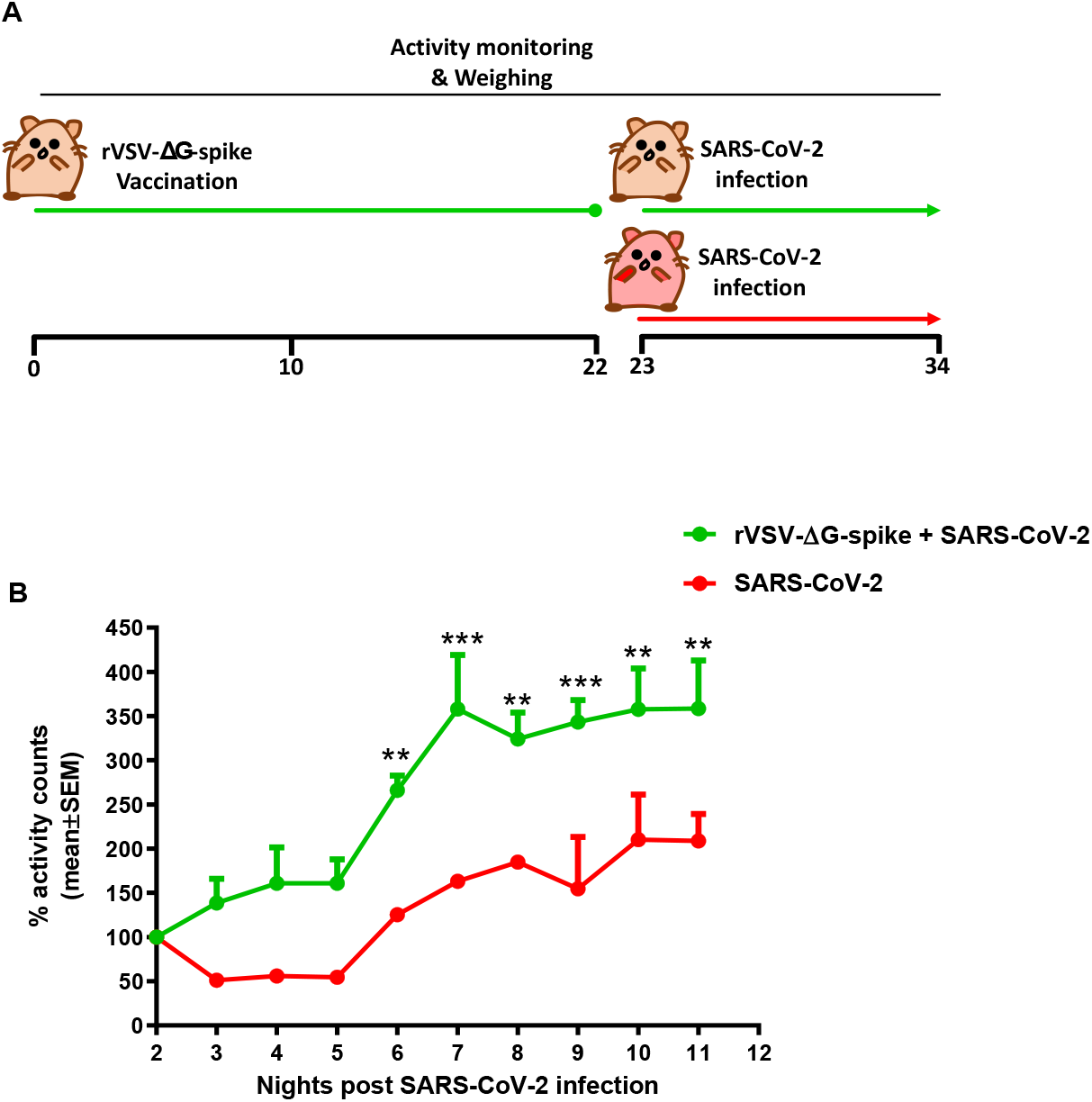
Recovery of group activity in females hamsters vaccinated with rVSV-ΔG-spike following SARS-CoV-2 infection. (A) rVSV-ΔG-spike vaccination and SARS-CoV-2 infection scheme. (B) Recovery of night time group activity of immunized hamsters compared to non-immunized, 2 nights after i.n. infection with 5×10^6^ pfu of SARS-CoV-2. Recovery was determined as percent of activity recorded at disease state 2 days after infection. n=2 cages/group, 4 hamsters/cage. **p<0.01, ***p<0.001

## Materials and methods

### Cell lines and viruses

African green monkey kidney clone E6 cells (Vero E6, ATCC^®^ CRL-1586™) were grown in Dulbecco’s modified Eagle’s medium (DMEM) containing 10% Fetal bovine serum (FBS), MEM non-essential amino acids (NEAA), 2mM L-Glutamine, 100Units/ml Penicillin, 0.1mg/ml Streptomycin, 12.5 Units/ml Nystatin (P/S/N) (Biological Industries, Israel). Cells were cultured at 37°C, 5% CO_2_ at 95% air atmosphere. For hamster’s infection, we used SARS-CoV-2 (GISAID accession EPI_ISL_406862), kindly provided by Bundeswehr Institute of Microbiology, Munich, Germany. For K18-hACE2 transgenic mice infection, we used SARS-CoV-2, isolate Human 2019-nCoV ex China strain BavPat1/2020 that was kindly provided by Prof. Dr. Christian Drosten (Charité, Berlin) through the European Virus Archive – Global (EVAg Ref-SKU: 026V-03883). Virus stocks were propagated and tittered on Vero E6 cells. The viruses were stored at −80°C until use. Handling and working with SARS-CoV-2 virus were conducted in a BSL3 facility in accordance with the biosafety guidelines of the Israel Institute for Biological Research (IIBR).

### Animal experiments

All animal experiments involving SARS-CoV-2 were conducted in a BSL3 facility. Treatment of animals was in accordance to Animal Welfare Act and the conditions specified in the Guide for Care and Use of Laboratory Animals (National Institute of Health, 2011). Animal studies were approved by the local IIBR ethical committee for animal experiments (protocols numbers HM-01-20, HM-02-20, and M-52-20). Female and male golden Syrian hamsters (60-90 gr., Charles River Laboratories, USA) 6-7 weeks old and female K18-hACE2 transgenic mice (18-20 gr., Jackson, USA) 8-10 weeks old, were maintained at 20−22°C and a relative humidity of 50 ± 10% on a 12 h light/dark cycle. Animals were fed with commercial rodent chow (Altromin, Germany) and provided with tap water ad libitum. Hamsters and mice were randomly assigned to experiment and kept in groups of 4 and 5, respectively. Both hamsters and mice were acclimated at their home cage for 3-5 days prior to infection to monitor and record the baseline of group activity.

Infection was performed as previously described^6^. Briefly, SARS-CoV-2 was diluted in PBS supplemented with 2% FBS (Biological Industries, Israel) and was used to infect anesthetized hamsters (5×10^6^ pfu) and mice (2000 pfu) by 50 μl and 20 μl respectively, by intranasal (i.n.) instillation of viral suspension.

Vaccination with rVSV-ΔG-spike was performed as previously described^6^. Briefly, intramuscular (i.m.) 50 μl/animal of rVSV-ΔG-spike (1×10^6^ pfu/animal) were administered to golden Syrian hamsters 23 days prior to i.n. infection with SARS-CoV-2.

### Animal housing and HCMS100 monitoring

For the home cage monitoring system - HCMS100, we used an industry standard home cage (Tecniplast^®^ 1285L home cage) and a non-limiting industry standard cage rack (Tecniplast^®^ DGM rack). Monitoring group activity of hamsters and mice in communal home cage by HCMS100 was performed as previously described7. Briefly, each cage is adjusted with a single retroflective laser sensor (HT3CL1/4P-M8, Leuze electronics, Germany) comprises of emitter and receiver, mounted adjacent to the home cage. Spontaneous movements of the rodents in the cages as indicated by a laser beam crosses situated in the drinking and feeding area, are continuously analyzed and recorded. All detectors are connected to communication controller and data collection and analyses are operated via remote user interface. Bin duration of laser crosses recording was every 10 minutes.

### Statistical analysis

Group activity counts are presented as means ± SEM. When noted, some group activity data are presented as mean ± SEM of percent of baseline activity of the same group. Weights are presented as mean ± SEM percent change from baseline of the same group. Group differences are analyzed by two ways ANOVA (group × time, with repeated measure on the latter) followed by an appropriate *post hoc* analysis (Dunnett) using GraphPad 7. A value of *p* < 0.05 was accepted as statistically significant.

## Discussion

The coronavirus disease 2019 (COVID-19) pandemic caused by SARS-CoV-2 infection has led to substantial unmet need for treatments. The development of such treatments will require testing them in an appropriate COVID-19 animal model. The pre-clinical research presented here centered on the detection of “disease state” in two animal species that can appropriately model the COVID-19 disease in humans, namely the Syrian golden hamsters and K18-hACE2 mice ^1^. COVID-19 disease in hamsters is often evaluated based on changes in their body weight, but is rarely evaluated by their overt symptoms. Thus, the only disease symptoms observed in SARS-CoV-2 infected hamsters were subtle with mild clinical features^3^. In this study we focused on changes in spontaneous group activity of hamsters in their home cage using the HCMS100, as an effective way to define and characterize changes in hamster’s health. We monitored group activity of hamsters in their home cage and described consistent pattern of high night time activity and low day time activity of these nocturnal animals with peak activity detected at about two hours into the night (Fig. 1). Similar pattern was obtained in hamsters in a single animal testing^8^ and this pattern is different from the two-peaks pattern at light shifts seen in groups of mice^7^.

Following intranasal exposure to SARS-CoV-2 we showed a significant decrease in overall group activity of hamsters both at the night time active phase and at the day time rest phase. This decrease started 1-2 days after infection. Male and female hamsters show similar pattern of disease state although females are generally more active than males during the night. This method of continuous monitoring of group activity was used together with the monitoring of weight loss as markers of disease progression and recovery. Comparisons between these two markers, group activity and weight loss, suggest that disease state can be detected as early as 48h post infection by the decrease in group activity that precedes the decrease in total body weight (Fig. 2). This may be explained by the notion that SARS-CoV-2 infection led to the reduced activity which may in turn decrease food consumption and induced weight loss. Weight loss seemed to be more prominent in males than in females^4^ while activity decreased more in females than in males^9^. These differences are probably the result of gender differences in baseline at this age: females are more active than males and males gain more weight than females. Thus, the use of weight loss as the sole marker of the disease, may lead to a possible interpretation bias.

K18-hACE2 transgenic mice expressing human angiotensin I converting enzyme 2 (ACE2)^5^ were recently established to model human COVID-19 disease and also serve as a reliable model for anti SARS-CoV-2 neutralizing antibodies treatment ^10–13^. We observed a sharp decrease in group activity and weight loss that was seen 5 days following viral exposure and death at 6-7 days post infection. Our data are consistent with the elegant study by Winkler et al., where assessment of COVID-19 disease progression in K18-hACE2 was evaluated using single animal treadmill stress test^11^. However, the use of the HCMS100 presented here eliminated the disruptions in mice habitat and social behavior and the stress associated with it^14^. In addition, HCMS100 remote, hands free method of observation better addressed the safety issues associated with the experimental use of SARS-CoV-2 virus in BSL3 facilities.

Convalescent hamsters previously exposed to SARS-CoV-2 and recovered are immune to SARS-CoV-2 and are expected to be protected from re-exposure to the same virus strain. This was also demonstrated here by monitoring group activity using the HCMS100 as well as the body weight changes. After the second exposure to SARS-CoV-2 (35 days after the first exposure), group activity was almost unaffected in convalescent hamsters, compared to naïve group that exhibited the expected decrease in activity post infection. This decrease in activity preceded the decrease in weight loss clearly seen in hamsters exposed to the virus for the first time. The weight of the newly infected animals returned to baseline level at approximately 20 days post infection while convalescent hamsters returned to their initial weight 5 days post infection. (Fig. 4D).

Continuous group activity monitoring was further used to study the efficacy of the newly developed rVSV-ΔG-spike vaccine^6^. In this pilot study, immunized animals were compared to a group of non-vaccinated hamsters in their response to SARS-CoV-2 infection. Following infection, both groups exhibited decreased night time activity. However, the group pre-immunized with rVSV-ΔG-spike vaccine showed a shorter morbidity phase as depicted by a faster recovery to baseline levels (Fig. 5).

The experiments described here demonstrate the importance of the use of behavioral monitoring of infected animals. Applying home cage monitoring system in studies of disease progression and treatments is on the rise^15^, however most available monitoring systems are limited to test a single animal per cage which enforces the untoward isolation of the experimental animals. In addition, the relatively high cost of these systems prohibits their use in large scale experiments commonly used in this type of research. The advantages of the HCMS100 used here was previously detailed^7^ and include the continuous monitoring of group activity for as long as required with minimal experimenters’ interference. This is a clear advantage in experimental designs that require exposure to pathogens such as the SARS-CoV-2 that restricts access to the experimental animals.

To date, this HCMS100 monitoring system was successfully used to detect changes in home cage continuous group activity of mice, hamsters and rats (unpublished data) with no need of alteration in either hardware or software of the HCMS100. Thus, the system can be easily applied to tests additional small animal species in the process of establishing pre-clinical evaluation of vaccines and drugs for human diseases.

